# Super-resolution modularity analysis shows polyhedral caveolin-1 oligomers combine to form scaffolds and caveolae

**DOI:** 10.1101/495382

**Authors:** Ismail M. Khater, Qian Liu, Keng C. Chou, Ghassan Hamarneh, Ivan Robert Nabi

**Author notes:** Correspondence (IRN), (GH). Equal contribution (Ghassan Hamarneh, Ivan Robert Nabi).

## Abstract

Caveolin-1 (Cav1), the coat protein for caveolae, also forms non-caveolar Cav1 scaffolds. Single molecule Cav1 super-resolution microscopy analysis previously identified caveolae and three distinct scaffold domains: smaller S1A and S2B scaffolds and larger hemispherical S2 scaffolds. Application here of network modularity analysis of SMLM data for endogenous Cav1 labeling in HeLa cells shows that small scaffolds combine to form larger scaffolds and caveolae. We find modules within Cav1 blobs by maximizing the intra-connectivity between Cav1 molecules within a module and minimizing the inter-connectivity between Cav1 molecules across modules, which is achieved via spectral decomposition of the localizations adjacency matrix. Features of modules are then matched with intact blobs to find the similarity between the module-blob pairs of group centers. Our results show that smaller S1A and S1B scaffolds are made up of small polygons, that S1B scaffolds correspond to S1A scaffold dimers and that caveolae and hemi-spherical S2 scaffolds are complex, modular structures formed from S1B and S1A scaffolds, respectively. Polyhedral interactions of Cav1 oligomers therefore leads progressively to the formation of larger and more complex scaffold domains and the biogenesis of caveolae.

## INTRODUCTION

Caveolae are smooth 50-80 nm plasma membrane invaginations whose formation requires the coat protein Cav1 and the adaptor protein CAVIN1 (also called PTRF) ^1^. Functional roles of caveolae include: mechanoprotective membrane buffers; mechanosensors; signaling hubs; and endocytic transporters ^2^. Cryo-electron microscopy (cryoEM) analysis of caveolae has reported that the Cav1 coat is polygonal, formed of distinct edges and suggested to form a dodecahedral cage ^3,4^. CryoEM analysis of Cav1 protein distribution in the caveolae coat, in either mammalian cells or following heterologous Cav1 expression in bacteria (h-caveolae), show that Cav1 exhibits a highly regular distribution of repeating polygons ^3,5,6^. CAVIN1 forms an outer filamentous coat layer whose filamentous structure likely corresponds to the striations observed on caveolae, as well as flattened caveolae, by deep-etch EM ^3,4,6–8^. In the absence of CAVIN1, Cav1 is localized to non-caveolar membrane domains known as Cav1 scaffolds ^9,10^. While scaffolds have been characterized functionally ^11,12^, defining their structure has proven more difficult. Biochemical analysis identifies small 8S oligomers that correspond to SDS-resistant oligomers of 10-15 Cav1 molecules as well as larger 60S oligomers that correspond to the caveolae coat ^13,14^. CryoEM suggests that small 8S oligomers combine to form the caveolar coat ^3,4^. However, the structural relationship of scaffolds to caveolae remains an open question. The size of both caveolae and scaffolds is below the diffraction limit of visible light (∼200-250 nm) and cannot be distinguished by diffraction limited microscopy.

Super-resolution microscopy is therefore ideally suited to identify and characterize these sub-diffraction limit cellular structures. Of the various super-resolution microscopy approaches, the best resolution is obtained using single-molecule localization microscopy (SMLM), based on the repeated activation (blinking) of small numbers of discrete fluorophores, such as *d*STORM, PALM, MINFLUX ^15–18^. In *d*STORM, precise localization of these blinks is determined from a Gaussian fit of the point spread function (PSF) providing ∼10-15 nm X-Y (lateral) resolution and ∼30 nm Z (axial) resolution for astigmatic lens 3D SMLM ^19,20^. SMLM generates point coordinates in 3D space that can then be used to reconstruct localizations with significantly improved resolutions and has been applied to distinguish invaginated and flattened caveolae based on Cav1 density in clusters ^21^. An alternate approach to study point distributions is to visualize them as a graph or network. Graphs are mathematical structures used to model interactions between entities for many systems, with the entities represented as graph nodes and the connections between them as edges ^22^. Real world graphs are frequently complex networks that have many different subgraphs or modules ^23^. Networks with high modularity have dense connections (edges) between the nodes within modules (sub-networks) and sparse connections between nodes in different modules. The optimization problem of finding divisions within a network (i.e. modules or communities) has been solved via various methods such as normalized-cut graph partitioning and spectral algorithms ^24,25^. Network and subgraph (module) analysis is therefore ideally suited to define molecular and subgroup organization between labeled molecules within the 3D SMLM point cloud of macromolecular complexes.

Previously, network analysis of SMLM Cav1 data sets in PC3 prostate cancer cells ^26^, that express Cav1 but no CAVIN1 and therefore no caveolae ^1^, identified two classes of Cav1 scaffolds corresponding to small Cav1 homo-oligomers (S1 scaffolds) that correspond to 8S Cav1 oligomers ^14,27^, as well as larger hemispherical S2 scaffolds. The formation of curved Cav1 structures in the absence of CAVIN1 is consistent with Cav1 induction of invaginated h-caveolae in bacteria and supports a role for Cav1 in membrane curvature ^5,28^. To identify signatures for caveolae, we compared PC3 prostate cancer cells that lack caveolae with PC3 cells transfected with the CAVIN1 adaptor required for caveolae formation. Larger hollow caveolae were only detected upon transfection of PC3 cells with CAVIN1 (PC3-PTRF cells) ^26^ and their modular nature supported the polyhedral Cav1 coat structure observed by cryoEM ^3,4^.

We now process (using spectral decomposition) the array of distances between localizations to find modules in endogenous Cav1 domains of HeLa cells. To determine the relationship between Cav1 scaffold domains and the multimeric caveolae structure, we leveraged a multi-threshold modularity analysis to extract the modules of the various Cav1 blobs. We then matched the blobs with the sub-modules of the various Cav1 domains based on the similarity (i.e., smaller Euclidean or L2 norm) between their biosignatures. To enhance localization precision, we used an SMLM microscope equipped with real-time nanometer-scale drift correction hardware ^29^ and included only high-precision localizations to improve localization accuracy ^30,31^. With this in-house built SMLM microscope, the localization precision approaches 10 nm ^32^, and the drift is limited to 1 nm in the x-y plane and 3 nm in the z axis ^29,33^. Modularity analysis and group matching show that S1A scaffolds can dimerize to form S1B scaffolds and oligomerize to form hemispherical S2 scaffolds. S1B scaffolds match the modules that make up the caveolae coat suggesting that the caveolae coat is built progressively by dimerization of S1A scaffolds, composed of the basic polygonal Cav1 units, that then combine to form a polyhedral caveolae coat.

## RESULTS AND DISCUSSION

### Tunable iterative merge algorithm

A major challenge to determining molecular structure by SMLM is defining molecular localizations (i.e. the location of the labeled molecule, in this case Cav1) from the millions of blinks (i.e. 3D spatial coordinates of the labelled Cav1 fluorescent events) generated from the stochastic blinking detected by SMLM. Many blinks derive from the same labeled molecule, particularly when the labeling approach is based on antibody labelling (i.e. *d*STORM). The same fluorophore can blink twice in succeeding acquisition frames dependent on the on-off duty cycle ^34^ and the same molecule can be labeled by different fluorophores, either on the same secondary antibody or on different secondaries bound to the same target protein, introducing error in blink localization relative to the actual antigen. Each of these blink localizations is in addition subject to localization error due to drift and to Gaussian fitting of the PSF. Multiple, distinct blink localizations therefore derive from the same molecule and generate a dense non-biological network (NBN) with high degree nodes centred around the actual molecule ^26,35^. Network analysis of the biological network, composed of nodes corresponding to predicted molecular localizations of the labeled proteins, requires reduction/consolidation of the NBNs.

Several methods have been proposed to reduce this artifact using temporal or spatial fluorophore information. Annibale et al. ^36,37^ proposed a method to correct the multiple-blinking in PALM by determining the merging time for mEos2 photoactivatable fluorescent protein. Other methods ^38–40^ spatially merged nearby localization events. Here, to correct for multiple-blinking and estimate molecular localization, we adopt the iterative merging algorithm of Khater et al. ^26^, which iteratively merges nearby nodes (blinks), that are within a threshold merging distance, until convergence is reached. The process starts with the high network degree nodes and continues until the distance between all pairs of reconstructed nodes, that correspond to predicted or estimated molecular localizations, is within the threshold merging distance. Nodes in closest proximity are combined first such that merging is initiated within the dense NBNs and continues progressively until no nodes within the point cloud are closer than the merging proximity threshold (MPT).

3D point clouds of Cav1 from 10 *d*STORM images of HeLa cells were processed using the *3D SMLM Network Analysis* computational pipeline ^26^. To address the multiple-blinking artifact that may bias the quantification process, we applied a tunable MPT from 10-20 nm in steps of 1 nm. Importantly, 4 classes were learnt at each MPT from 10-20 nm. Further, tuning the MPT from 10-20 nm minimally impacted classification, size, modularity, characteristic path and hollowness of all 4 classes of blobs (Fig. 1A). Machine learning blob classification is therefore independent of the merge algorithm for MPTs from 10-20 nm. Not unexpectedly, increasing the MPT reduced the predicted molecular localization number per blob. We set the MPT based on the reported 145 Cav1 proteins per caveolae ^41^. An MPT of 19 nm resulted in an average of 142 localizations for the largest H2 blobs (Fig. 1A), that match the PP2 caveolae blobs from PC3-PTRF cells (see Fig. 2A).

**Figure 1.**
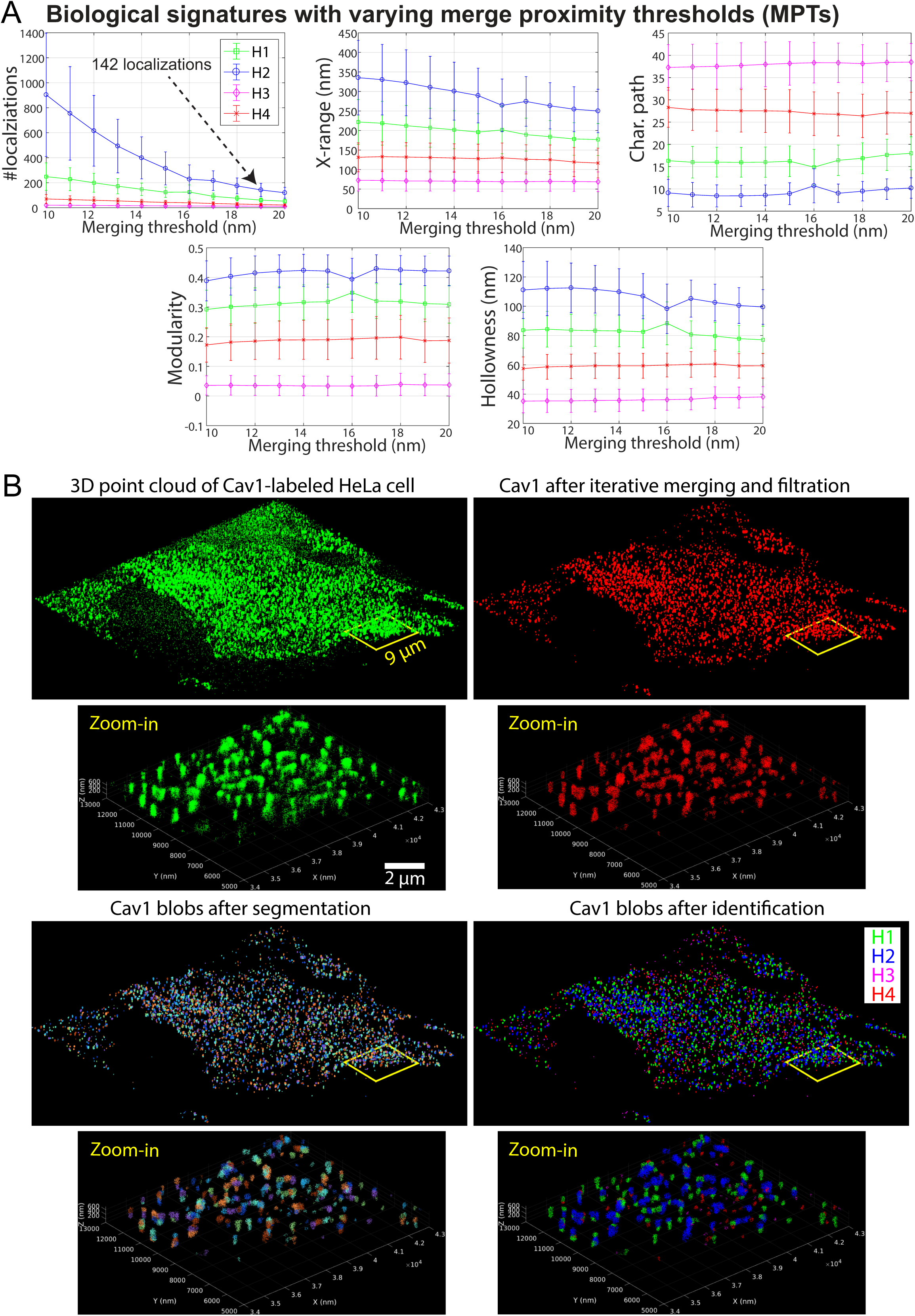
MPT tuning does not impact blob identification. **A.** Biological signatures of HeLa Cav1 blobs at different MPTs (10-20 nm) were obtained by *3D SMLM Network Analysis* ^26^. We learn 4 groups/classes of Cav1 domains at each MPT. Cav1 blob shape, topology, hollowness and network features are minimally affected by MPT tuning while number of molecular localizations is affected by MPT tuning. Error bars represent standard deviation. **B.** 3D Cav1 point clouds of a representative HeLa cell imaged with drift-corrected *d*STORM ^29,33^ before (green) and after (red) iterative blink merging at 19 nm and filtering out noisy localizations. Color-coded representations of blobs after segmentation and after identification by machine learning using *3D SMLM Network Analysis* ^26^ pipeline are shown. We identified four groups of blobs representing different Cav1 domains in HeLa cells.

**Figure 2.**
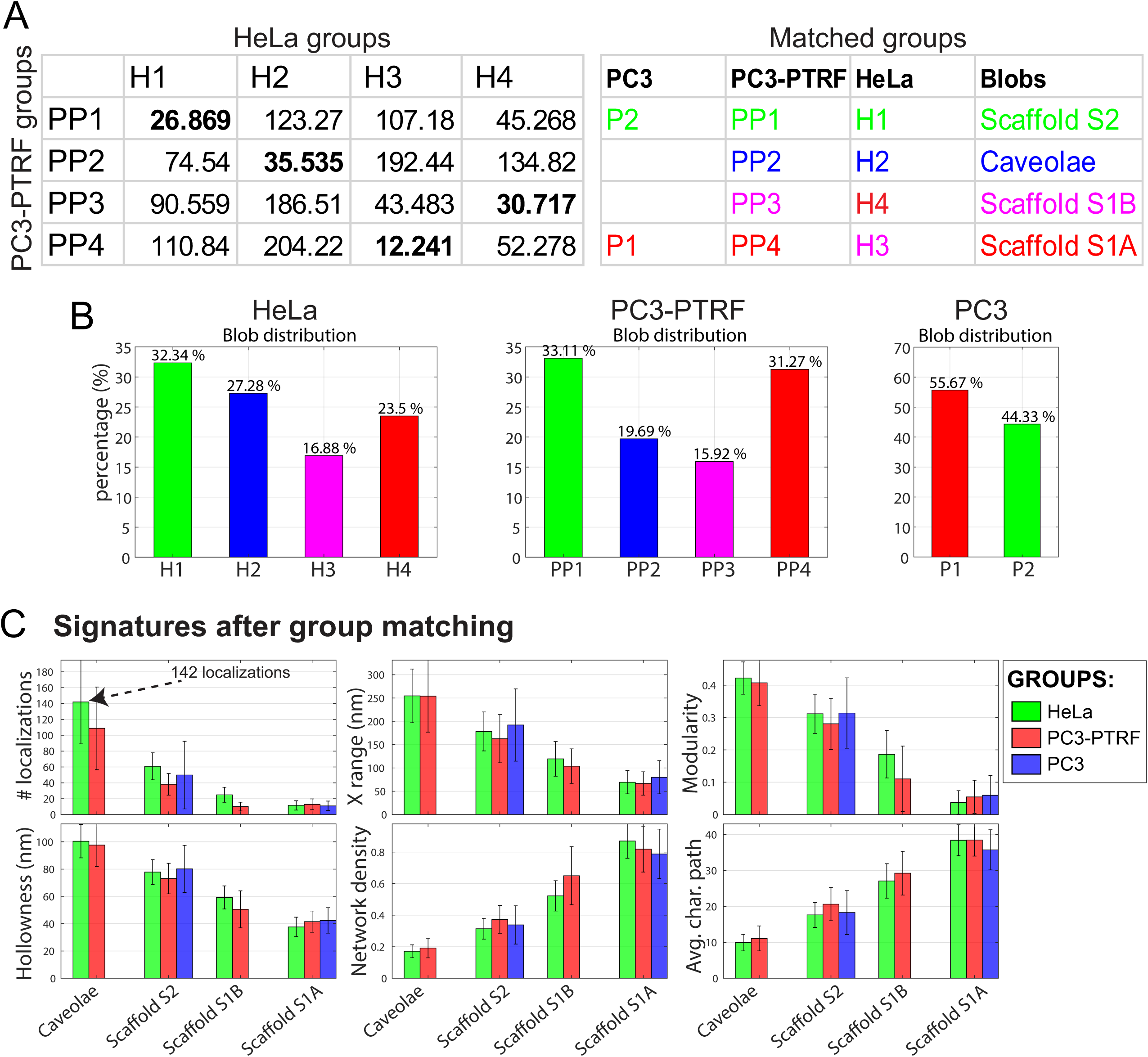
*3D SMLM Network Analysis* of the HeLa cells dataset. **A.** Matching HeLa Cav1 groups with previously identified Cav1 domains in PC3 and PC3-PTRF cells ^26^. The numbers are the Euclidean distances that capture the similarity/dissimilarity between the groups with smaller numbers indicating increased similarity. We matched learned groups from PC3, PC3-PTRF and HeLa cells and show distances among the feature vector of group centers (in bold are the closest matching groups). The table to the right shows color matching of HeLa groups with previously identified P1 and P2 Cav1 domains in PC3 cells and PP1, PP2, PP3, and PP4 Cav1 domains in PC3-PTRF cells ^26^. **B.** Distribution of the matched groups from HeLa, PC3-PTRF and PC3 datasets are presented for comparison. **C.** Signatures of matched groups from HeLa (at 19 nm MPT), PC3 and PC3-PTRF (at 20 nm MPT) cells show a high degree of correspondence of the individual group features. See Supp. Fig. S1 for the rest of the features.

Figure 1B shows the 3D point cloud of one of the HeLa cells in our dataset at various stages of the pipeline: 1) The 3D point cloud of Cav1-labeled HeLa cell generated by real-time drift control SMLM; 2) After iterative merging and denoising filtration. The denoising module visits every Cav1 event and predicts whether it is signal or noise. This prediction is based on examining the network features for every Cav1 event in our data, as well as examining corresponding network features of nodes in a random network. If the network features of a Cav1 event are similar to those of the random network’s nodes, then that Cav1 event will be declared as noise and removed. This denoising process will retain the Cav1 clusters (blobs) and filter out noisy localizations and monomeric Cav1; 3) After segmentation into separate blobs and extraction of a 28 feature/descriptor vector for every blob; 4) After unsupervised machine learning to learn the various Cav1 domains from the extracted blobs and their descriptor features.

### Group matching

Machine learning identified four groups of Cav1 domains (H1, H2, H3, and H4) in HeLa cells (Fig. 1A). We used the Euclidean distance in 28 dimensions to encode similarity of HeLa groups with groups previously identified in PC3 and PC3-PTRF cells ^26^, with similarity proportional to the inverse Euclidean distance. As seen in Figure 2A, for the groups with larger blobs, H2 matches PP2, corresponding to caveolae, and H1 matches PP1, corresponding to the larger hemispherical S2 scaffolds. For the smaller S1 scaffolds, H4 matches PP3 and H3 matches PP4. Distribution of the different classes of blobs in the different cell types shows that HeLa and PC3-PTRF cells present a similar distribution of Cav1 blobs with slightly more caveolae detected in HeLa cells (Fig. 2B).

Feature analysis after group matching shows that the four HeLa groups match with high degree the four PC3-PTRF groups as well as the S2 and S1A scaffolds present in PC3 cells (Fig. 2C; see also Supp. Fig. S1 for additional data on blob features). Relative to the PC3 data ^26^, we observed a doubling in molecular localizations for S1B scaffolds relative to S1A scaffolds and increased modularity of S1B scaffolds in the HeLa data set that we attribute to the improved resolution obtained with the real-time drift control SMLM ^29,33^. Increased Cav1 localization number in S1B scaffolds parallels the increased size (X-range) and reduced network density of these clusters relative to S1A scaffolds, reflecting differences between these structures that led to their classification as distinct cluster groups in this and our previous analysis ^26^. Indeed, we observe a progressive increased number of localizations (Caveolae > S2 Scaffolds > S1B scaffolds > S1A scaffolds) associated with increased modularity and decreased network density (Fig. 2C). Caveolae are the most modular structures (modularity > 0.4), then S2, and S1B. S1A scaffolds have the least tendency to form modules (modularity < 0.04).

### Small Cav1 S1 scaffolds combine to build larger scaffolds and caveolae

The variable modularity of the different classes of Cav1 blobs led us to extract the blobs’ modules and study their features. To make sure that we correctly construct the modules within the blob’s network, we used multi-proximity threshold (PT) network analysis (Fig. 3A) to decompose the blobs’ networks into modules using spectral analysis. For all the groups, any PT greater than 60 nm renders every blob a single connected component; the average connected component size plateaus and equals the blob size for the different groups at PTs greater than 60 nm. This indicates that at PT > 60 nm, all post-merge Cav1 localizations in a cluster are within 60 nm of each other. The number of modules and of Cav1 localizations per module are stable across the PT range from 60 to 170 nm. This range is therefore suitable to determine the number of modules and of Cav1 localizations per module. HeLa caveolae were found to be highly modular containing 6-7 modules of ∼29 Cav1 localizations, S2 scaffolds 5 modules of ∼14 localizations and S1B scaffolds ∼4 modules of 7-8 localizations each (Fig. 3A). S1A scaffolds have the minimum average number of modules of ∼2 modules per blobs of ∼5-6 localizations each.

**Figure 3.**
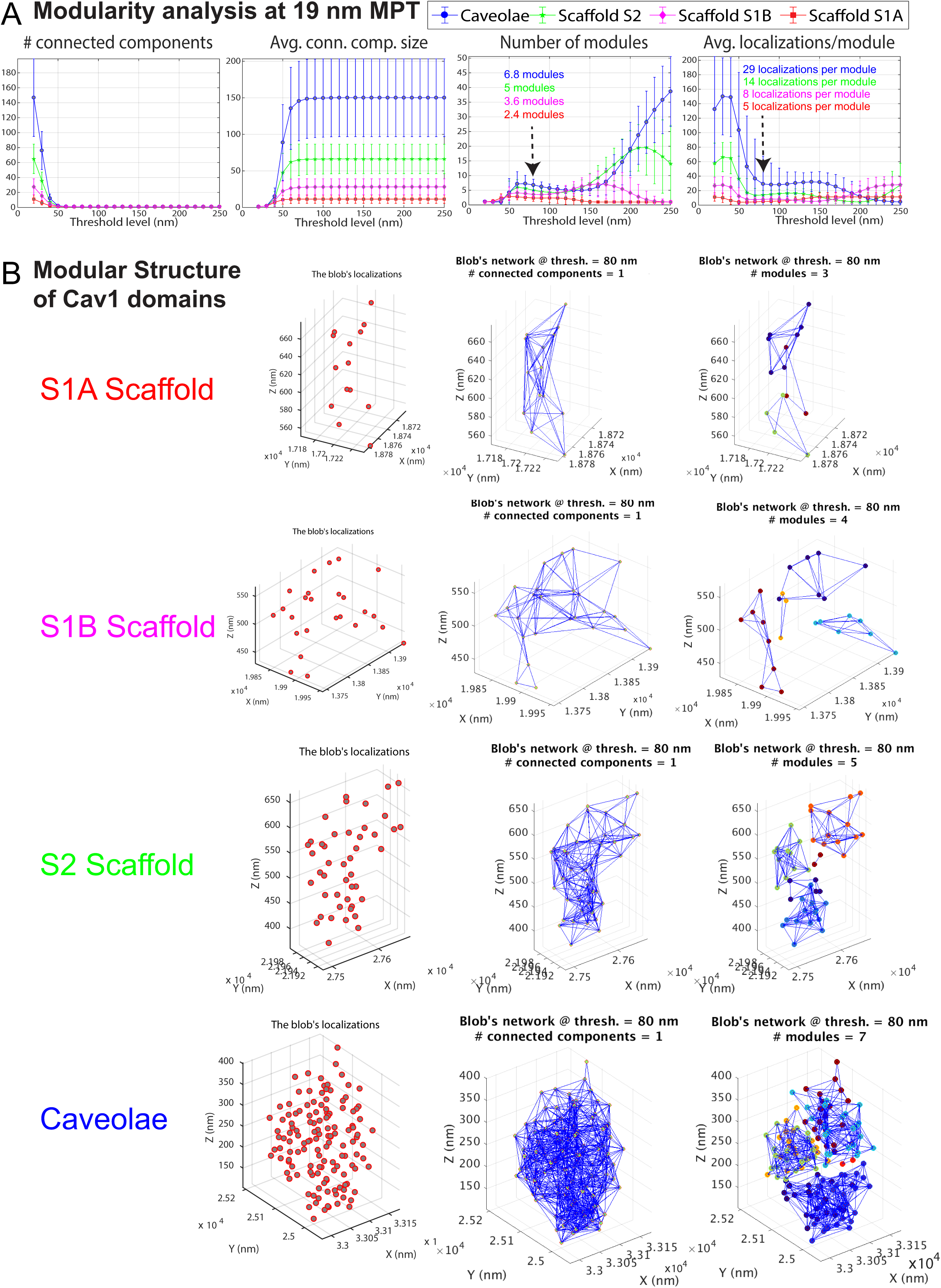
Modularity analysis of Cav1 blobs. **A.** Multi-proximity threshold modularity analysis shows the number of connected components, number of modules and localizations per module (at 19 nm MPT) for HeLa blobs at different proximity thresholds. **B.** Representative blobs from the different HeLa Cav1 domains are shown. Visualization shows the blob’s localizations, the localizations’ connections, and the blob’s modules.

Visualization of blobs from the identified groups (Fig. 3B) highlights the modular nature of the various Cav1 structures. At 80 nm, each blob forms one connected component network and extracted modules for every blob are shown in different colors. The presence of small modules (∼5-8 molecules) within both S1A and S1B scaffolds is indicative of an additional degree of suborganization within these small scaffold domains. 3D cryoEM tomography identified a network of 3-way junctions and polygonal arrangements of Cav1 protein densities within the caveolae coat ^3^. Similarly, cryoEM analysis of Cav1-induced vesicles in bacteria (h-caveolae) present distinct polygonal repeating units on the h-caveolae cage ^5^. We propose that the sub-modules that we detect in S1A and S1B scaffolds correspond to these polygonal repeating units that comprise the caveolae coat. The fact that S1A scaffolds form one connected component unit and that the number of localizations of S1A scaffolds matches that of Cav1 homo-oligomers (∼14-15 Cav1s) ^14,27^ suggests that interaction between these polygonal sub-modules forms more stable structural units. This is supported by the identification of larger modules in both S2 scaffolds and caveolae (Fig. 3A,4A).

**Figure 4.**
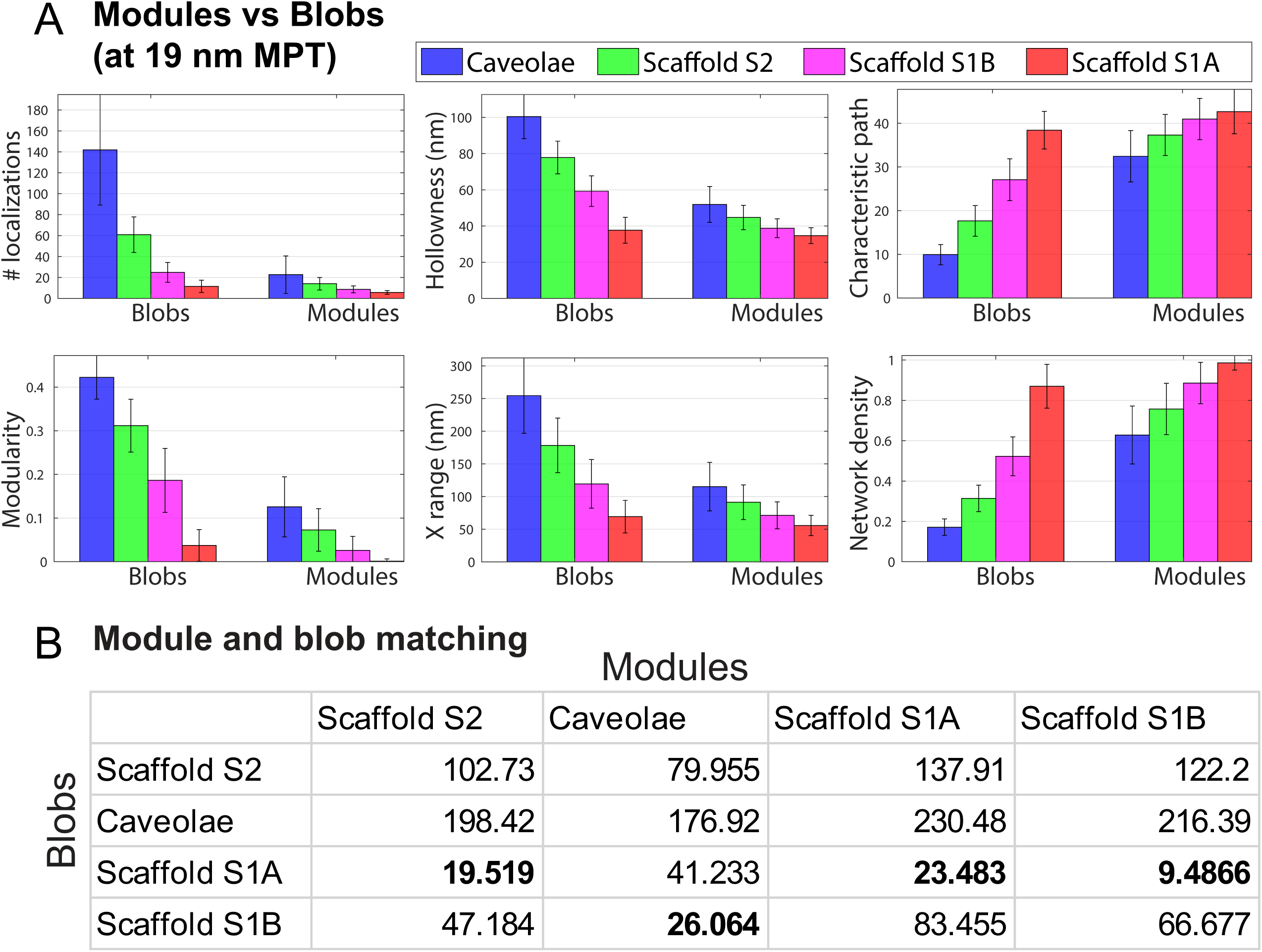
Module-blob matching between Cav1 domains. **A.** Signatures of Cav1 blobs and blob modules shows that some module features are similar to blob features. For example, the right bars that represent the caveolae modules (blue) are very similar to the left bars that represent the S1B blobs (magenta). **B.** We extracted 28 features (e.g. shape, topology, hollowness, network) for every blob and module. The table encodes the module-blob similarity between the different Cav1 domains (blobs) and the modules of each type as Euclidean distances between every pair of group centres.

Most interestingly, the decomposed modules from the different Cav1 cluster groups show a much higher degree of similarity in terms of the shape, topology and network features than the clusters from which they originate (Fig. 4A). For instance, while Cav1 blob classes show a progressive reduction in network density from S1A scaffolds to caveolae, modules from the different blob classes show a similar network density. This suggests that differential interaction between modules is responsible for the changes in network density of the different classes of Cav1 blobs and that these modules form fundamental building blocks of larger Cav1 structures.

Indeed, many features of caveolae modules match S1B scaffolds while S2 and S1B modules match S1A scaffolds. For instance, #localizations, hollowness, characteristic path, modularity, size, and network density of the caveolae modules (blue bars to right of graphs) are very similar to their corresponding features in the S1B blobs (magenta bars to left) (Fig. 4A). We quantitatively assessed module-blob similarity across all features using the matching matrix of the features for the various blobs and modules group center using Euclidean distance (Fig. 4B). The column-wise (i.e. the modules) similarity shows that: S2 scaffold modules match S1A blobs; caveolae modules match S1B blobs; and S1B modules match S1A blobs. The close matching of S1B modules with S1A blobs and doubling in number of modules and localizations of S1B modules relative to S1A blobs suggests that S1B scaffolds represent dimers of S1A scaffolds. Further, PC3 cells that lack caveolae have only S1A and S2 scaffolds (Fig. 2 B,C) ^26^ supporting the matching between S1A blobs and S2 modules reported here. The dissimilarity between caveolae and S2 scaffolds and the modules of any other blob types suggests that these are complex structures made up of primitive S1A and S1B scaffolds.

Overall, our data support a model in which Cav1 is organized into smaller units of 5-8 Cav1 localizations that correspond to the polygonal base units observed by cryoEM analysis of the Cav1 caveolar coat ^3,5^. These base units combine to form larger stable structures of which the smallest is S1A scaffolds, that we propose correspond to the previously identified ∼14-15 Cav1 homo-oligomers ^14,27^. We also identify S1B scaffolds, previously classified as distinct from S1A scaffolds ^26^ as larger structures that may correspond to S1A dimers. Modularity analysis and group matching show that S1A scaffolds combine to form both S1B dimers and the larger hemispherical S2 scaffold structures. Caveolae modules show better matching and correspond in size to S1B and not S1A modules suggesting that caveolae formation may be a two-step process in which S1A scaffolds first combine to form dimers that then interact to form the caveolae coat (Fig. 5, Video S1). Consistent with a role for S1B in caveolae formation, PC3 cells that lack caveolae and CAVIN1 do not contain S1B scaffolds ^26^. As cluster size increases, Cav1 domains show a more pronounced reduction in density relative to their constituent modules. This suggests that interaction between smaller S1 scaffolds to form larger structures, including caveolae, is associated with changes in how modules interact and are organized. Importantly, our analysis based on TIRF microscopy argues that all 4 Cav1 domains, from S1A scaffolds to caveolae are present at the plasma membrane. Based on the role of caveolae as membrane buffers that flatten in response to mechanical stretching ^42^, we suggest that these modular interactions are dynamic and reversible.

**Figure 5.**
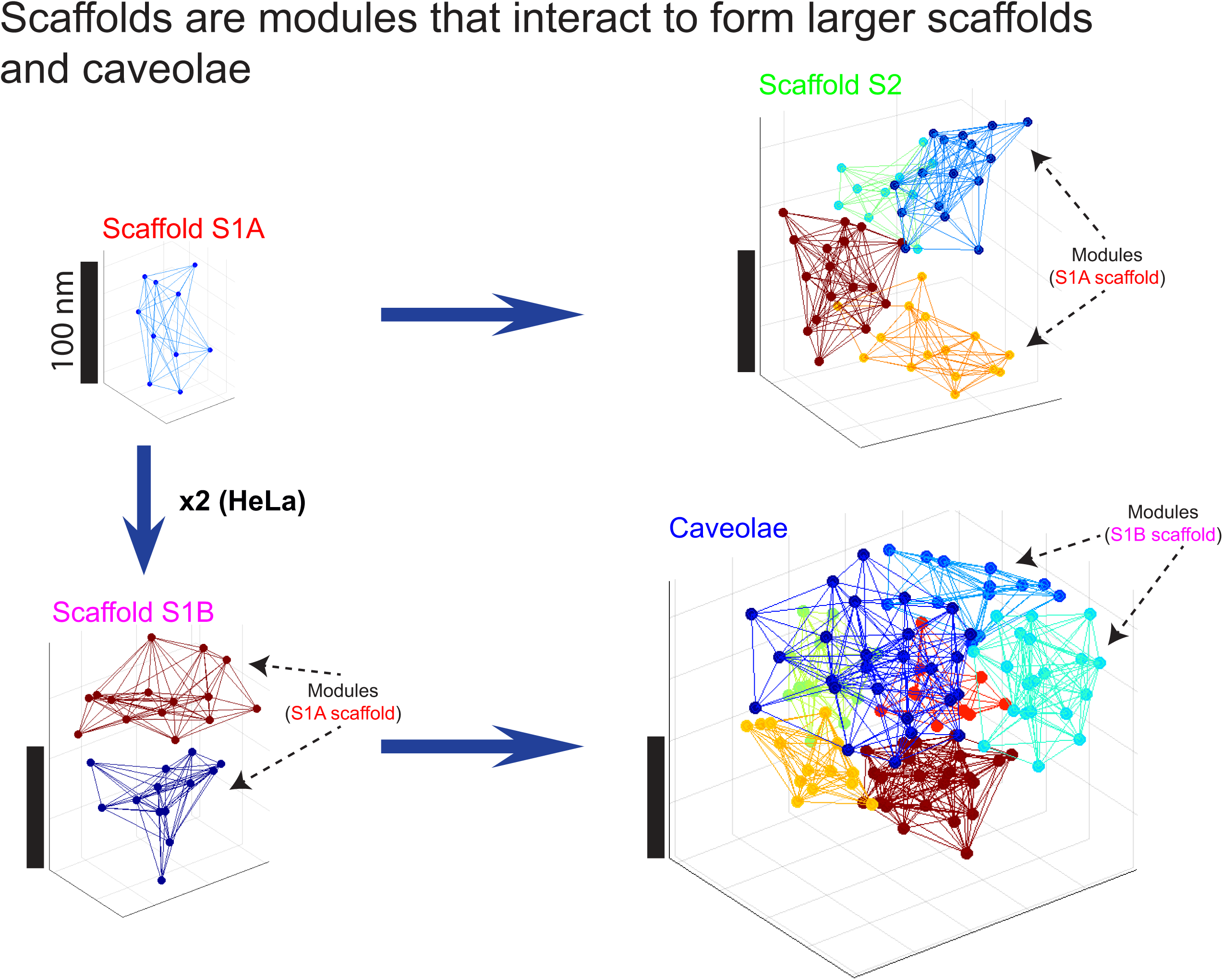
Modular interaction of Cav1 S1A scaffolds forms larger scaffolds and caveolae. Based on the module-blob matching results (Fig. 4B), S1A blobs are stable primitive structures (simplex) that are used to build up more complex, modular S1B and S2 scaffolds. S1B scaffolds correspond to S1A dimers and are used to build the caveolae coat complex (see Video S1). The figure also shows the hemispherical shape of S2 blobs and the hollow caveolae blobs.

In this work, we applied multi-threshold modularity analysis to networks/blobs constructed from 3D point clouds of Cav1 localizations acquired via SMLM. Classification of endogenous Cav1 domains in HeLa cells matched those previously identified in CAVIN1-transfected PC3 prostate cancer cells ^26^. Spectral decomposition allowed us to define the relationship between the different Cav1 blobs and their extracted sub-networks/modules via biosignature similarity and matching across the various domains/groups. This approach is applicable for modular analysis of other oligomeric macromolecular biological structures improving our understanding of their architecture by disassembling them into their basic building components.

## MATERIALS AND METHODS

### Cell culture and immunofluorescent labeling

HeLa cells were tested for mycoplasma by PCR (Applied Biomaterial, Vancouver, BC, Canada) and cultured in Dulbecco’s Modified Eagle’s medium (DMEM; Invitrogen) containing 10% fetal bovine serum (Invitrogen). For SMLM imaging, cells were plated on fibronectin coated coverslips (No. 1.5H) for 24 h prior to fixation with 3% paraformaldehyde (PFA) for 15 min at room temperature. Fixed cells were rinsed with PBS/CM (phosphate buffered saline complemented with 1 mM MgCl_2_ and 0.1 mM CaCl_2_), permeabilized with 0.2% Triton X-100 in PBS/CM, and blocked with 10% goat serum and 1% bovine serum albumin (BSA; Sigma-Aldrich Inc.) in PBS/CM before incubation with rabbit anti-caveolin-1 (BD Transduction Inc.) for 12 h at 4°C and then Alexa Fluor 647-conjugated goat anti-rabbit (Thermo-Fisher Scientific Inc.) for 1 h at room temperature. Primary and secondary antibodies, at saturating concentrations of antibodies, were diluted in SSC (saline sodium citrate) buffer containing 1% BSA, 2% goat serum and 0.05% Triton X-100. Cells were washed extensively after each antibody incubation with SSC buffer containing 0.05% Triton X-100 and post-fixed using 3% PFA for 15 min followed by extensive washing with PBS/CM. Near-infrared fiducial markers (diameter 100 nm; Thermo Fisher Scientific) were added for real-time drift correction. Immediately prior to imaging, cells were mounted and sealed on glass depression slides in freshly prepared imaging buffer (10% glucose, 0.5 mg/ml glucose oxidase, 40 μg/mL catalase, 50 mM Tris, 10 mM NaCl and 50 mM β-mercaptoethylamine (Sigma-Aldrich Inc.) in double-distilled water ^20,34^.

### SMLM Imaging

Imaging of Hela cells was performed on an in-house built SMLM system equipped with an apochromatic TIRF oil immersion objective lens (60x/1.49; Nikon Instruments) and a real-time drift correction system which limits the lateral drift to ∼1 nm and the axial drift to ∼3 nm. A 639 nm laser line (Genesis MX639, Coherent Inc., USA) was used to excite Alexa Fluor 647 fluorophores and near-infrared fiducial markers. A 405 nm laser line (Laserglow Technologies) was used to activate Alexa Fluor 647. The detailed optical setup and the imaging acquisition procedure were described previously ^29,33^.

The dataset used in this work consists of 10 fields of view (FOV) of Cav1-labeled HeLa cells. Each HeLa FOV is 54×54×1 μm^3^ which is 9 times larger than the FOV in the PC3 cell study (18×18×1 μm^3^), acquired using a Leica GSDIM microscope equipped with a 160X objective ^26^. Each HeLa FOV therefore included multiple cells and this study analysed a larger number of cells than the PC3 study. We collected 40,000 frames per super resolution image. The total number of collected localizations that we processed per image ranged from 1.6 to 6.7 million. The *3D SMLM Network Analysis* method was able to process the whole FOV. Details of the *3D SMLM Network Analysis* approach can be found in ^26^.

### Multi-proximity threshold network modularity analysis

For every blob of 3D localizations, more than one network can be constructed; one per each proximity threshold in the set {PT_1_, PT_2_,…, PT_T_} (i.e. *blob_i_* has *T* networks 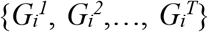, where 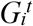 is composed of a set of nodes *V_i_* and edges 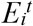 to form 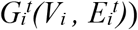. *V_i_*, unaffected by PT_*t*_, represents the molecules of *blob_i_* and 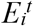 is the set of edges connecting all pairs of molecules interacting within PT_*t*_ nm.

We leverage a spectral decomposition algorithm to find modules within Cav1 blobs. Given 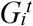, a *blob_i_*’s network at PT_*t*_, we find its modules (communities) using the Newman method ^24,44^. Specifically, we first calculate an adjacency matrix whose element *m, n* encodes the distance between the m-*th* and *n-th* localizations. A spectral decomposition method calculates the eigenvector representation of this adjacency matrix. This eigendecomposition defines the modules as it maximizes the intra-connectivity between Cav1 molecules within a module and minimizes inter-connectivity between Cav1 molecules across modules.

Given 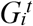, a *blob_i_*’s network at PT_*t*_, we find the optimal number of modules (communities) using eigenvectors of the network adjacency matrix. At small PTs, the molecules of a blob might not form one connected network (i.e. the network might consist of more than one connected component). A blob network containing non-dense and non-connected regions cannot be used to extract modules (i.e. as per definition, networks with high modularity have dense connection between the nodes within modules and sparse connections between nodes in different modules). Hence, PTs that generate networks with more than one connected component should be avoided when extracting the modules.

### Features extraction and module-blob similarity

For every segmented Cav1 blob, we extracted 28 features that are then used to group the blobs’ into classes using the *3D SMLM Network Analysis* pipeline ^26^. The learned classes from HeLa dataset are then matched with the previously identified Cav1 domains from PC3 and PC3-PTRF datasets ^26^. The modules of the HeLa Cav1 blobs are then extracted using the multi-proximity threshold network modularity analysis described in the previous subsection. We extracted 28 features for every module. To find the similarity/dissimilarity among the extracted modules and the various blobs, we leveraged the matching analysis to match blob modules with intact blobs using the Euclidean distance of group centers.

## ACKNOWLEDGMENTS

Supported by grants from the CIHR (PJT-156424, PJT-159845), NSERC and CFI/BCKDF (IRN, GH, KCC).

## AUTHOR CONTRIBUTIONS

Conceptualization, I.M.K., I.R.N. and G.H.; Methodology, I.M.K., I.R.N. and G.H; Software, I.M.K. and G.H.; Sample Preparation, Q. L.; SMLM Imaging, Q. L. and K. C. C.; Formal Analysis, I.M.K., I.R.N. and G.H.; Investigation, I.M.K., I.R.N. and G.H.; Resources, K.C.C., I.R.N. and G.H.; Writing – Original Draft, I.M.K., I.R.N. and G.H.; Writing – Review & Editing, I.M.K., I.R.N. and G.H.; Visualization, I.M.K., I.R.N. and G.H.; Supervision, I.R.N. and G.H.; Project Administration, I.R.N. and G.H.; Funding Acquisition, K.C.C., I.R.N. and G.H.

---

**Supp. Figure S1.**
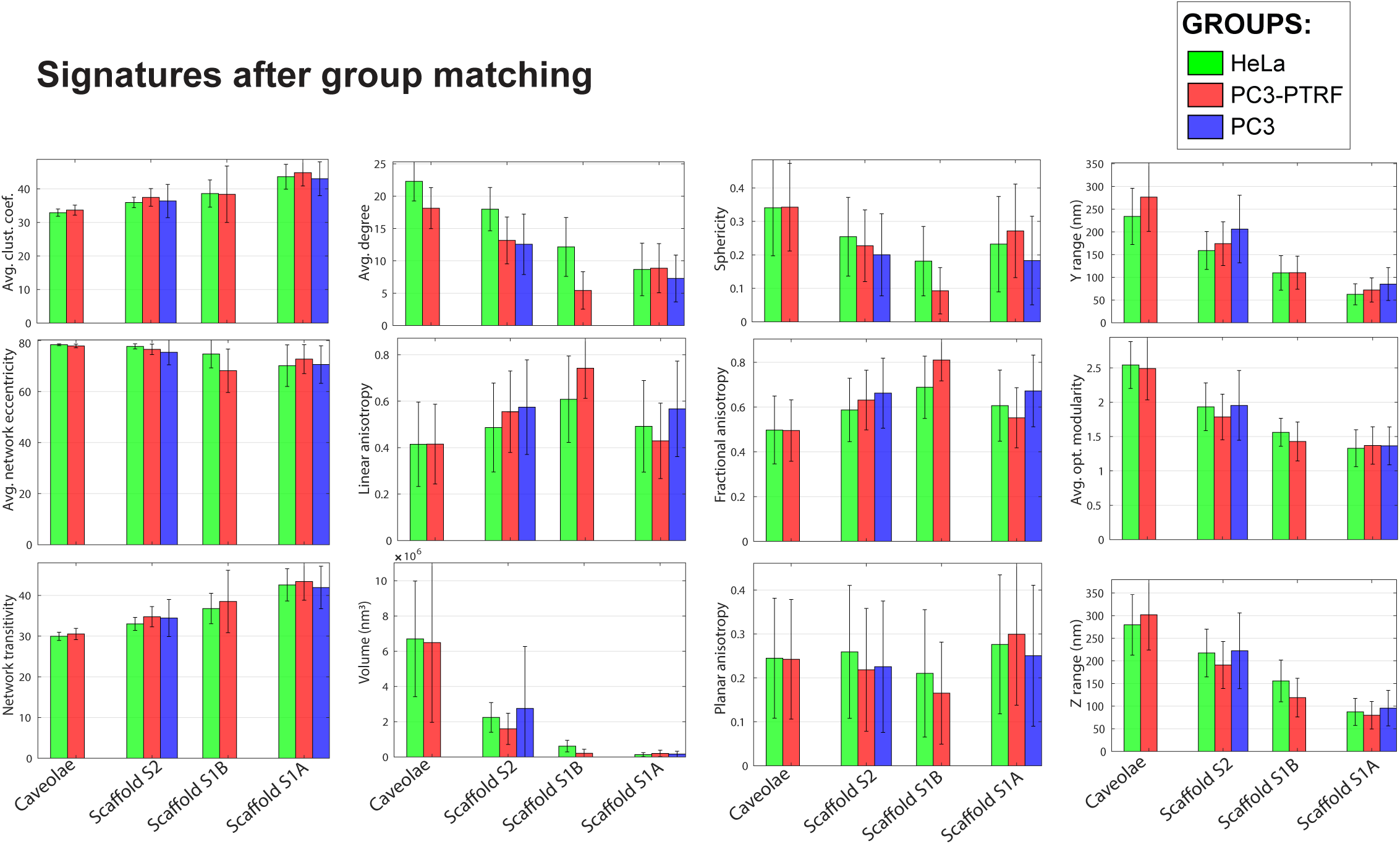
Additional features for Cav1 blobs. Cav1 blob biosignatures in addition to those presented in Fig. 2C are shown for the matched groups from PC3, CAVIN1/PTRF-transfected PC3 (PC3-PTRF), and HeLa cells

**Supp. Video S1.** Animated rotating structures of S1A scaffolds (H3), S1B scaffolds (H4), S2 scaffolds (H1), and caveolae (H2) blobs in HeLa cells. The blob’s localizations, connections between localizations (edges), and modules (for S1B, S2 scaffolds and caveolae) are shown from different angles with rotation. The video illustrates Cav1 domain formation as described in Figure 5. Rotating blobs in 3D enhances visualization of the internal structure of the blobs. For example, the hollow nature of caveolae is clear when rotating the views.

## REFERENCES

1 Hill, M. M. et al. PTRF-Cavin, a conserved cytoplasmic protein required for caveola formation and function. Cell 132, 113–124 (2008).

2 Parton, R. G. & del Pozo, M. A. Caveolae as plasma membrane sensors, protectors and organizers. Nat. Rev. Mol. Cell Biol. 14, 98–112 (2013).

3 Ludwig, A., Nichols, B. J. & Sandin, S. Architecture of the caveolar coat complex. J. Cell Sci. 129, 3077–3083, doi:10.1242/jcs.191262 (2016).

4 Stoeber, M. et al. Model for the architecture of caveolae based on a flexible, net-like assembly of Cavin1 and Caveolin discs. Proc. Natl. Acad. Sci. USA 113, E8069–E8078, doi:10.1073/pnas.1616838113 (2016).

5 Walser, P. J. et al. Constitutive formation of caveolae in a bacterium. Cell 150, 752–763, doi:10.1016/j.cell.2012.06.042 (2012).

6 Ludwig, A. et al. Molecular composition and ultrastructure of the caveolar coat complex. PLoS bBiology 11, e1001640 (2013).

7 Kovtun, O. et al. Structural insights into the organization of the cavin membrane coat complex. Dev. Cell 31, 405–419, doi:10.1016/j.devcel.2014.10.002 (2014).

8 Rothberg, K. G. et al. Caveolin, a protein component of caveolae membrane coats. Cell 68, 673–682 (1992).

9 Head, B. P. & Insel, P. A. Do caveolins regulate cells by actions outside of caveolae? Trends Cell Biol. 17, 51–57 (2007).

10 Lajoie, P., Goetz, J. G., Dennis, J. W. & Nabi, I. R. Lattices, rafts, and scaffolds: domain regulation of receptor signaling at the plasma membrane. J. Cell Biol. 185, 381–385 (2009).

11 Lajoie, P. et al. Plasma membrane domain organization regulates EGFR signaling in tumor cells. J. Cell Biol. 179, 341–356 (2007).

12 Moon, H. et al. PTRF/cavin-1 neutralizes non-caveolar caveolin-1 microdomains in prostate cancer. Oncogene 33, 3561–3570, doi:10.1038/onc.2013.315 (2014).

13 Hayer, A., Stoeber, M., Bissig, C. & Helenius, A. Biogenesis of caveolae: stepwise assembly of large caveolin and cavin complexes. Traffic 11, 361–382, doi:10.1111/j.1600-0854.2009.01023.x (2010).

14 Monier, S. et al. VIP21-caveolin, a membrane protein constituent of the caveolar coat, oligomerizes in vivo and in vitro. Mol. Biol. Cell 6, 911–927 (1995).

15 Rust, M. J., Bates, M. & Zhuang, X. Sub-diffraction-limit imaging by stochastic optical reconstruction microscopy (STORM). Nature Methods 3, 793–795, doi:10.1038/nmeth929 (2006).

16 Eilers, Y., Ta, H., Gwosch, K. C., Balzarotti, F. & Hell, S. W. MINFLUX monitors rapid molecular jumps with superior spatiotemporal resolution. Proc. Natl. Acad. Sci. USA 115, 6117–6122, doi:10.1073/pnas.1801672115 (2018).

17 Betzig, E. et al. Imaging intracellular fluorescent proteins at nanometer resolution. Science 313, 1642–1645, doi:10.1126/science.1127344 (2006).

18 Balzarotti, F. et al. Nanometer resolution imaging and tracking of fluorescent molecules with minimal photon fluxes. Science 355, 606–612, doi:10.1126/science.aak9913 (2017).

19 Shroff, H., Galbraith, C. G., Galbraith, J. A. & Betzig, E. Live-cell photoactivated localization microscopy of nanoscale adhesion dynamics. Nat Methods 5, 417–423, doi:http://www.nature.com/nmeth/journal/v5/n5/suppinfo/nmeth.1202_S1.html (2008).

20 Huang, B., Wang, W., Bates, M. & Zhuang, X. Three-Dimensional Super-Resolution Imaging by Stochastic Optical Reconstruction Microscopy. Science 319, 810–813, doi:10.1126/science.1153529 (2008).

21 Tachikawa, M. et al. Measurement of caveolin-1 densities in the cell membrane for quantification of caveolar deformation after exposure to hypotonic membrane tension. Sci Rep 7, 7794, doi:10.1038/s41598-017-08259-5 (2017).

22 Newman, M. E. J. The structure and function of complex networks. SIAM review 45, 167–256 (2003).

23 Kim, J. & Wilhelm, T. What is a complex graph? Physica A: Statistical Mechanics and its Applications 387, 2637–2652 (2008).

24 Newman, M. E. J. Modularity and community structure in networks. Proc. Natl. Acad. Sci. USA 103, 8577–8582 (2006).

25 Newman, M. E. J. Spectral methods for community detection and graph partitioning. Physical Review E 88, 042822 (2013).

26 Khater, I. M., Meng, F., Wong, T. H., Nabi, I. R. & Hamarneh, G. Super Resolution Network Analysis Defines the Molecular Architecture of Caveolae and Caveolin-1 Scaffolds. Sci Rep 8, 9009, doi:10.1038/s41598-018-27216-4 (2018).

27 Sargiacomo, M. et al. Oligomeric structure of caveolin: implications for caveolae membrane organization. Proc. Natl. Acad. Sci. USA 92, 9407–9411 (1995).

28 Ariotti, N. et al. Molecular Characterization of Caveolin-induced Membrane Curvature. J. Biol. Chem. 290, 24875–24890, doi:10.1074/jbc.M115.644336 (2015).

29 Tafteh, R. et al. Real-time 3D stabilization of a super-resolution microscope using an electrically tunable lens. Opt. Express 24, 22959–22970 (2016).

30 Foi, A., Trimeche, M., Katkovnik, V. & Egiazarian, K. Practical Poissonian-Gaussian noise modeling and fitting for single-image raw-data. IEEE Trans Image Process 17, 1737–1754, doi:10.1109/TIP.2008.2001399 (2008).

31 Thompson, R. E., Larson, D. R. & Webb, W. W. Precise nanometer localization analysis for individual fluorescent probes. Biophys. J. 82, 2775–2783, doi:10.1016/S0006-3495(02)75618-X (2002).

32 Liu, Q., Chen, L., Aguilar, H. C. & Chou, K. C. A stochastic assembly model for Nipah virus revealed by super-resolution microscopy. Nature Communications 9, 3050, doi:10.1038/s41467-018-05480-2 (2018).

33 Tafteh, R., Scriven, D. R., Moore, E. D. & Chou, K. C. Single molecule localization deep within thick cells; a novel super-resolution microscope. J Biophotonics 9, 155–160, doi:10.1002/jbio.201500140 (2016).

34 Dempsey, G. T., Vaughan, J. C., Chen, K. H., Bates, M. & Zhuang, X. Evaluation of fluorophores for optimal performance in localization-based super-resolution imaging. Nature Methods 8, 1027–1036, doi:10.1038/nmeth.1768 (2011).

35 Khater, I. M., Liu, Y, Liu, Q, Meng, F, Chou, KC, Nabi, IR, Hamarneh, G. Distinguishing biological and non-biological networks in single molecule localization super-resolution micrscopy. in American Society for Cell Biology and European Molecular Biology Organization (ASCB-EMBO) Vol. 28 page 1094 (Abstract P2728) (Molecular Biology of the Cell Philadelphia, USA, 2017).

36 Annibale, P., Vanni, S., Scarselli, M., Rothlisberger, U. & Radenovic, A. Quantitative photo activated localization microscopy: unraveling the effects of photoblinking. PloS one 6, e22678 (2011).

37 Annibale, P., Vanni, S., Scarselli, M., Rothlisberger, U. & Radenovic, A. Identification of clustering artifacts in photoactivated localization microscopy. Nature methods 8, 527 (2011).

38 Nieuwenhuizen, R. P. J. et al. Quantitative localization microscopy: effects of photophysics and labeling stoichiometry. PloS one 10, e0127989 (2015).

39 Andronov, L. et al. 3DClusterViSu: 3D clustering analysis of super-resolution microscopy data by 3D Voronoi tessellations. Bioinformatics 34, 3004–3012 (2018).

40 Andronov, L., Lutz, Y., Vonesch, J.-L. & Klaholz, B. P. SharpViSu: integrated analysis and segmentation of super-resolution microscopy data. Bioinformatics 32, 2239–2241 (2016).

41 Pelkmans, L. & Zerial, M. Kinase-regulated quantal assemblies and kiss-and-run recycling of caveolae. Nature (Lond.) 436, 128–133 (2005).

42 Sinha, B. et al. Cells respond to mechanical stress by rapid disassembly of caveolae. Cell 144, 402–413 (2010).

43 Newman, M. E. J. Finding community structure in networks using the eigenvectors of matrices. Physical review E 74, 036104 (2006).

